# integRATE: a desirability-based data integration framework for the prioritization of candidate genes across heterogeneous omics and its application to preterm birth

**DOI:** 10.1101/302612

**Authors:** Haley R. Eidem, Jacob Steenwyk, Jennifer Wisecaver, John A. Capra, Patrick Abbot, Antonis Rokas

**Affiliations:** Department of Biological Sciences, Vanderbilt University, Nashville, TN, USA; Department of Biochemistry, Purdue University, West Lafayette, IN, USA; Department of Biomedical Informatics, Vanderbilt University, Nashville, TN, USA; Vanderbilt Genetics Institute, Vanderbilt University, Nashville, TN, USA

**Keywords:** prematurity, integrative genomics, complex disease, candidate gene ranking, Venn diagram

## Abstract

**Background:** The integration of high-quality, genome-wide analyses offers a robust approach to elucidating genetic factors involved in complex human diseases. Even though several methods exist to integrate heterogeneous omics data, most biologists still manually select candidate genes by examining the intersection of lists of candidates stemming from analyses of different types of omics data that have been generated by imposing hard (strict) thresholds on quantitative variables, such as P-values and fold changes, increasing the chance of missing potentially important candidates.

**Methods:** To better facilitate the unbiased integration of heterogeneous omics data collected from diverse platforms and samples, we propose a desirability function framework for identifying candidate genes with strong evidence across data types as targets for follow-up functional analysis. Our approach is targeted towards disease systems with sparse, heterogeneous omics data, so we tested it on one such pathology: spontaneous preterm birth (sPTB).

**Results:** We developed the software integRATE, which uses desirability functions to rank genes both within and across studies, identifying well-supported candidate genes according to the cumulative weight of biological evidence rather than based on imposition of hard thresholds of key variables. Integrating 10 sPTB omics studies identified both genes in pathways previously suspected to be involved in sPTB as well as novel genes never before linked to this syndrome. integRATE is available as an R package on GitHub (https://github.com/haleyeidem/integRATE).

**Conclusions:** Desirability-based data integration is a solution most applicable in biological research areas where omics data is especially heterogeneous and sparse, allowing for the prioritization of candidate genes that can be used to inform more targeted downstream functional analyses.

## Background

Biological processes underlying disease pathogenesis typically involve a complex, dynamic, and interconnected system of molecular and environmental factors [1]. Advances in high-throughput omics technologies have allowed for the collection of data corresponding to the genomic, transcriptomic, epigenomic, proteomic, and metabolomic elements that contribute to variation in these biological processes [2]. However, each of these omics approaches, when employed in isolation, can only capture variation within a single layer of a much more complicated biological system [3,4]. For example, even though the thousands of single nucleotide polymorphisms (SNPs) that have been linked to complex diseases or traits via genome-wide association studies (GWAS) have greatly contributed to our understanding of complex disease, we still lack in depth knowledge of the molecular mechanisms underlying the vast majority of these associations [5]. Similarly, transcriptomics studies routinely identify hundreds to thousands of differentially expressed genes between diseased and healthy tissue samples, but disentangling the disease-causing changes in gene expression from its byproducts can be far more challenging [6]. Given the limitations of each omics approach and their focuses on different layers of the biological system, integration of different types of omics data to identify the key biological pathways involved in disease has emerged as a promising avenue for research [4].

One integrative study design is to obtain diverse types of omics data from the same tissue samples or patient cohorts. The resulting data can then be vertically integrated (Fig. 1A, top left) to identify candidate genes and pathways involved in complex disease. Alternatively, a single type of omics data can be collected from a variety of tissue samples or patient cohorts, facilitating their horizontal integration across many samples, which can substantially increase the experiment’s power (Fig. 1A, top right). In both vertical and horizontal integration study designs, the availability of diverse types of omics data from the same samples enables the use of a variety of statistical integration approaches (Fig. 1A, bottom) [7]. For example, multi-staged integration uses multiple steps to first identify associations between different data types and then identify associations between data types and the phenotype of interest [8], whereas meta-dimensional integration combines data simultaneously based on concatenation, transformation, or model building [9].

**Fig. 1.**
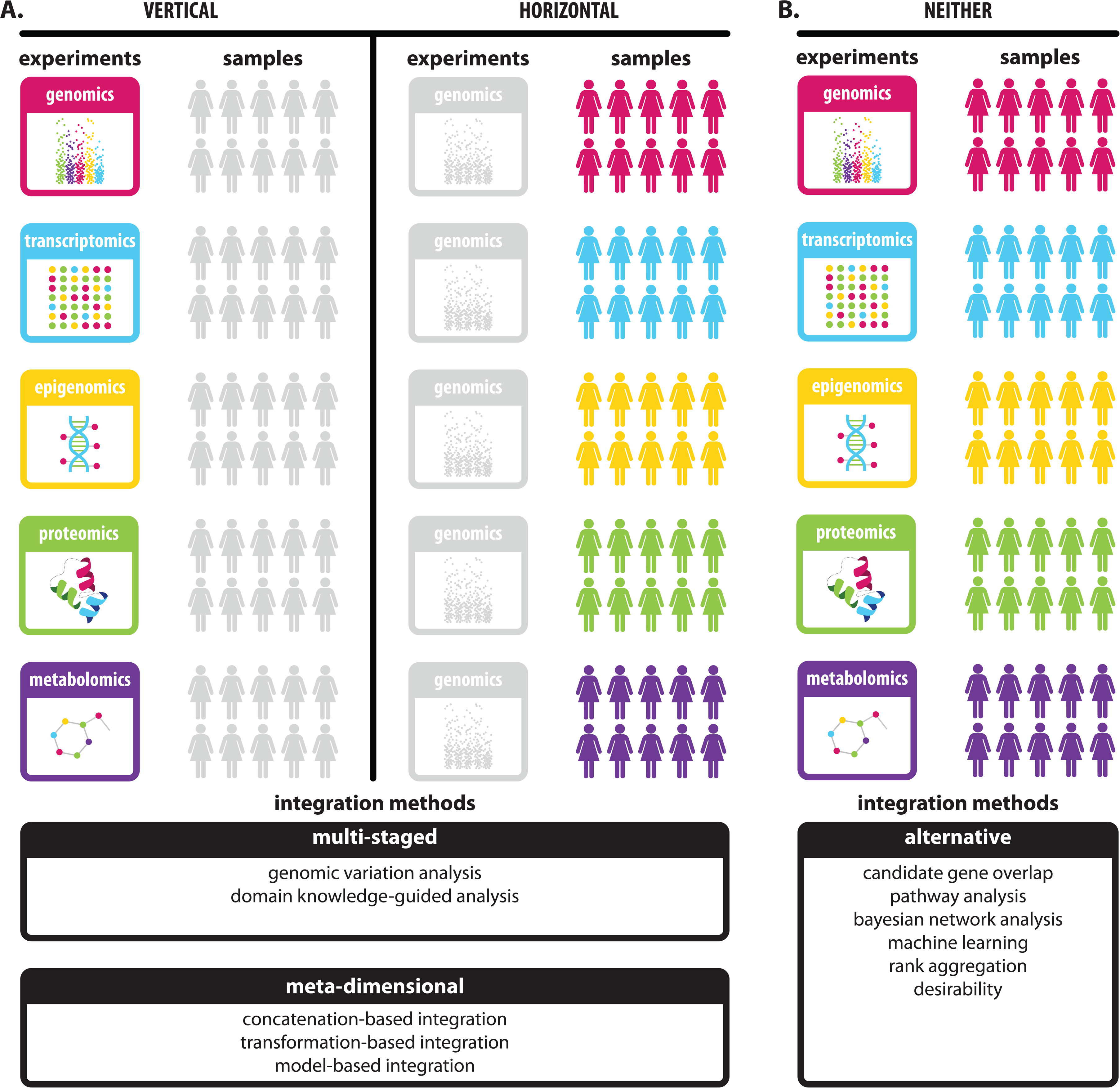
Selecting a data integration strategy depends on the structure of accessible multi-omics data. (A, left) If multiple types of omics data (‘multi-omics’) are available for the same cohort of patients, vertical integrative analysis can be performed to combine information across data types. This integration can be achieved using a variety of multi-staged and metadimensional statistical approaches that identify disease subtypes, regulatory networks, and driver genes. (A, right) If the opposite is true and a specific type of omics data is available across a number of different patient cohorts, horizontal meta-analysis can be performed to increase statistical power and identify disease-associated perturbations. (B) In some cases, however, experimental data are only available for different omics data types from different cohorts of patients and neither vertical nor horizontal data integration can be performed. In these situations, integration relies on mapping data to common units (e.g., genes or pathways) and then either integrating transformed data or simply overlapping candidate sets. The software approach presented here (integRATE) utilizes desirability functions to transform and integrate heterogeneous data allowing for the prioritization of candidate genes for functional analysis.

Although multi-omics data sets generated using vertical and horizontal study designs are becoming increasingly common, such data sets are lacking for many complex diseases [10-14]. Often, heterogeneous omics data are collected study by study, for a limited set of tissue samples and across only one or two omics data types at a time (Fig. 1B, top). For each study, a long list of genes or genomic regions with associated data is produced and sorted based on effect size (e.g., fold change), significance (e.g., P-value), or some other criterion. Hard thresholds can then be imposed on P-values, for example, to bin the genes or genomic regions and identify significant candidates for further analysis; this type of approach can then be applied across multiple, heterogeneous omics studies.

Several problems exist with the imposition of hard thresholds, however. Including (or excluding) genes or genomic regions as candidates based on P-value, fold change, expression level, and/or odds ratio cutoffs introduces biases and removes information, especially when combining multiple cutoffs from several criteria [15-17]. These cutoffs can sometimes even be arbitrary, like selecting the top n or n% from each data set. Additionally, statistical significance is not always equivalent to biological significance, meaning that non-statistically significant genes may still be involved in disease pathogenesis, or vice versa. Moreover, while selecting the top n genes might limit the scope of further functional analysis, the alternative approach of selecting all significant hits could mean that thousands of genes are identified as candidates. A final consideration in analyzing heterogeneous omics data is that we sometimes do not know any genes, pathways, or networks that have already been shown to be involved in complex disease. Some integration methods, especially those based on prediction (e.g., machine learning, network analysis), depend on the availability of such knowledge for algorithm training and cannot be performed in their absence [7,8,18-21].

Desirability functions provide a way to integrate heterogeneous omics data in systems where gold standards (i.e., genes known to be involved in the complex disease under investigation) are not yet known (Fig. 1B, bottom). Originally developed for industrial quality control, desirability functions have been successfully used in chemoinformatics to rank compounds for drug discovery and have been proposed as a way to integrate multiple selection criteria in functional genomics experiments [22-26]. In the context of integrating diverse but heterogeneous omics data, desirability functions allow for the ranking and prioritizing of candidate genes based on cumulative evidence across data types and their variables, rather than within-study separation of significant and non-significant genes based on single variables in single studies. For example, a 2015 study initially proposed the use of desirability functions to integrate multiple selection criteria for ranking, selecting, and prioritizing genes across heterogeneous biological analyses and demonstrated its use by analyzing a set of microarray-generated gene expression data [22].

To facilitate data integration in the presence of heterogeneous multi-omics data and when prior biological knowledge is limited, we propose a desirability-based framework to prioritize candidate genes for functional analysis. To facilitate application of our framework, we built a user-friendly software package called integRATE, which takes as input data sets from any omics experiment and generates a single desirability score based on all available information. This approach is targeted towards biological processes or diseases with particularly sparse or heterogeneous data, so we test integRATE on a set of 10 omics data sets related to spontaneous preterm birth (sPTB), a complex disease where heterogeneous multi-omics data are the only omics data currently available.

## Design

### Variable Transformation

First, relevant studies need to be identified for integration; this selection can be based on any number of characteristics including tissue(s) sampled, disease subtype, or experimental designs (Fig. 2, step 1). The data in each of these studies (e.g., gene expression data, proteomic data, GWAS data, etc.) are typically specific to or can be mapped to individual genetic elements (e.g., genes) in the genome. Furthermore, each study’s data contain genetic element-specific values for many different variables (e.g., P-value, odds ratio, fold change, etc.). Then desirability functions are fit to the observations for each variable within a study (e.g., P-value, odds ratio, fold change, etc.) according to whether low values are most desirable (*d_low_*, e.g., P-value), high values are most desirable (*d_high_*, e.g., odds ratio), or extreme values are most desirable (*d_extreme_*, e.g., fold change) (Fig. 2, step 2). The desirability score for each genetic element can be calculated by applying one of the following equations to a given variable:

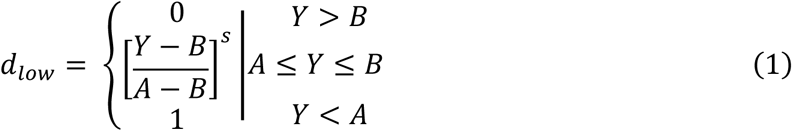

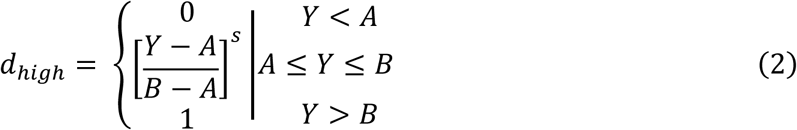

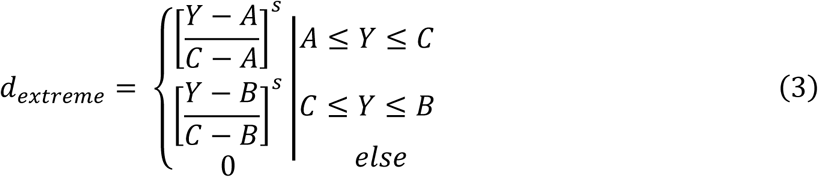

In these equations, *Y* is the variable value and *s* is the scale coefficient affecting the function’s rate of change that can be customized according to user preference. Alternatively, the equations could be used without any scaling by setting the scale coefficient to 1. For *d_low_* and *d_high_*, *A* is the low cut point and *B* is the high cut point where the function changes. For *d_extreme_*, *A* is the low cut point, *C* is the intermediate cut point, and *B* is the high cut point where the function changes. The user can customize these cut points based on numerical values (e.g., P-value < 0.05) or percentile values (e.g., top 10%). The resulting values, ranging from 0 to 1 (or the minimum and maximum values specified) are transformed desirability scores based on information from each variable.

**Fig. 2.**
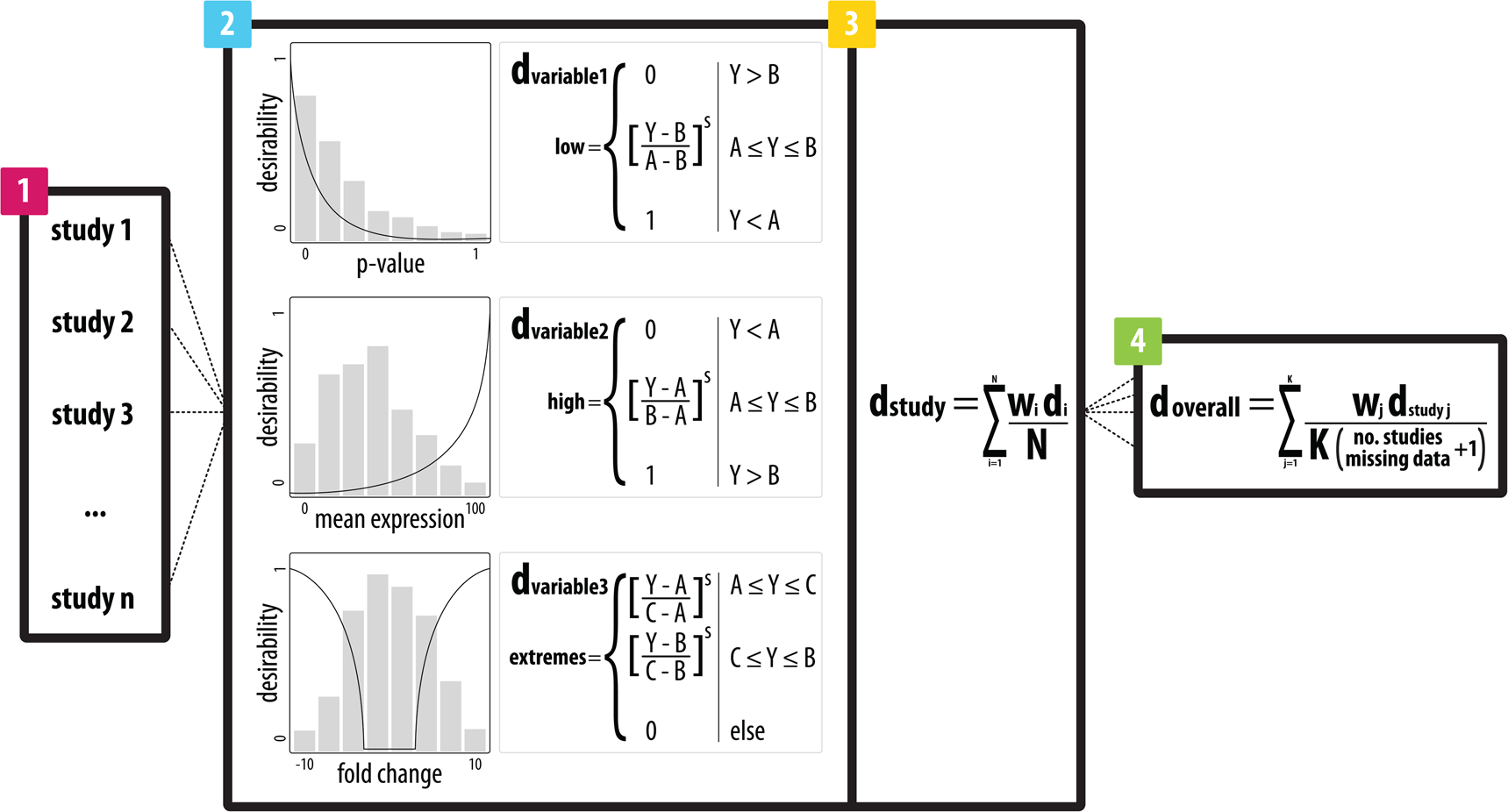
integRATE relies on three main steps to identify studies, integrate data, and rank candidate genes. (1) Relevant studies must first be identified for integration based on any number of features including, but not limited to: phenotype, experimental design, and data availability. (2) Data corresponding to all variables in each study are then transformed according to the appropriate desirability function. In this step, the user assigns a function based on whether low values are most desirable (*d_low_*), high values are most desirable (*d_high_*), or extreme values are most desirable (*d_extreme_*) and can customize the shape of the function with other variables like cut points (*A*, *B*, *C*), scales (*s*), and weights (*w*) to better reflect the data distributions or to align with user opinion regarding data quality and relevance. (3) These variable-based scores are integrated (*d_study_*) with a straightforward arithmetic mean (where weights can also be applied) to produce a single desirability score for each gene in each study containing information from all variables simultaneously. (4) Finally, study-based desirability scores are integrated to produce a single desirability score for each gene (*d_overall_*) that includes information from all variables in all studies and reflects its cumulative weight of evidence from each data set identified in step 1. These scores are normalized by the number of studies containing data for each gene and can be used to rank and prioritize candidate genes for follow up computational and, most importantly, functional analyses.

### Variable Integration

Next, desirability scores for each of the *N* variables within a study (e.g., P-value, odds ratio, fold change, etc.) are combined using an arithmetic mean so that genetic elements (e.g., genes) with desirability scores of zero for any given variable remain in the analysis (Fig. 2, step 3). Desirability for genetic elements within a study can be calculated by:

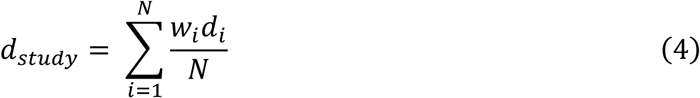

In this equation, *w_i_* is the weight parameter (assigned to each variable), *d_i_* is desirability score for each genetic element based on the values of each variable derived from equations (1), (2) or (3), and *N* is the total number of transformed variables. This step produces a single desirability score (*d_study_*) for each genetic element in the study containing information from all transformed variables. Here, the user is also able to include variable weights (*w_i_*) when integrating their desirability scores, which can be useful in cases where certain variables are considered more informative or accurate than others.

### Study Integration

Finally, the *d_study_* values can be integrated using the arithmetic mean to produce a single desirability score (*d_overall_*) for each genetic element, representing its desirability as a candidate according to the weight of evidence from all variables in all *K* studies that were integrated (Fig. 2, step 4). The overall score used to prioritize candidates can be calculated by:

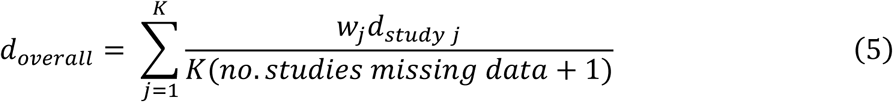

In this equation, *w_j_* is the weight parameter (assigned to each study), *d_study j_* is the desirability score for each study, and *K* is the total number of studies integrated. Importantly, the overall desirability score *d_overall_* is normalized by the number of studies missing data for each genetic element to account for the number of values contributing to each overall desirability score. This normalization factor can be used to calculate a soft cutoff for the most desirable candidates that is equivalent or higher than the desirability score that would be achieved by a genetic element with a perfect desirability score of 1 in a single study but missing from all other studies. We call genetic elements achieving desirability scores equal to or above this cutoff ‘desirable.’

### Software

The methodology described above is implemented in our software, integRATE, available on GitHub as an R package (https://github.com/haleyeidem/integRATE). Although we focus on using desirability functions to integrate heterogeneous omics data corresponding to complex human diseases, integRATE can be applied to data sets from any phenotype, species, and data type (provided that the units can all be mapped to a common set of elements, such as genes). Functionality is provided for the application of customizable desirability functions as well as data visualization.

## Implementation

One human complex genetic disease where the omics data available are heterogeneous is preterm birth (PTB). Defined as birth before 37 weeks of completed gestation, PTB is the leading cause of newborn death worldwide [27]. Although 30% of preterm births are medically indicated due to complications including preeclampsia (PE) or intrauterine growth restriction (IUGR), the remaining 70% occur spontaneously either due to the preterm premature rupture of membranes (PPROM) or idiopathically (sPTB). Further complicating factors are that multiple maternal and fetal tissues are involved (e.g., placenta, fetal membranes, umbilical cord, myometrium, decidua, etc.) as well as multiple genomes (maternal, paternal, and fetal) [28]. Evidence from family, twin, and case-control studies suggests that genetics plays a role in determining birth timing and a recent GWAS identified a handful of genes linked to prematurity [29]. Nevertheless, the pathogenesis of PTB and its many subtypes remains poorly understood [30-32].

The publicly available data for sPTB consist of several different independently conducted omics analyses that would be challenging to analyze with statistical approaches developed for vertical and horizontal integration [29,33,34]. Although these omics data have been analyzed in isolation, integration of their information using the desirability-based platform implemented in integRATE may provide unique insights into the complex mechanisms involved in regulating birth timing and, thus, allow for the identification and prioritization of novel candidate genes for further functional and targeted analyses.

### Study Identification

Studies were initially identified based on the following PubMed search strategy (10/19/2017):

> “Pregnancy”[mh] AND “Humans”[mh] AND “Preterm birth”[mh] AND (“Gene Expression Profiling”[mh] OR “Gene Expression Regulation”[mh]) AND (“Placenta”[mh] OR “Decidua”[mh] OR “Myometrium”[mh] OR “Cervix Uteri”[mh] OR “Extraembryonic Membranes”[mh] OR “Blood”[mh] OR “Plasma”[mh] OR “Umbilical Cord”[mh])

Studies that reported conducting a genome-wide omics analysis of sPTB from a preliminary scan of the abstract were downloaded for full-text assessment. Furthermore, a thorough investigation was conducted of their associated reference lists to identify studies not captured via PubMed.

Additionally, each study had to meet the following inclusion criteria:

1. Experimental group consisted of sPTB cases only and was not confounded by other pregnancy phenotypes (e.g., preeclampsia),
2. Analysis was genome-wide and not targeted to any specific subset of genes or pathways, and
3. Full data set was publicly available (not just top n%).

We identified 54 studies through the first phase of our literature search, but only 10 data sets that met all inclusion criteria. All excluded studies are listed in Additional File 1 with reasons for exclusion and the 10 data sets used in our pilot analysis are outlined in Table 1 [33-46].

**Table 1.** The 10 sPTB omics data sets identified for desirability-based integration.

| First Author | Year | Experiment | Control | Tissue | omics Type |
| --- | --- | --- | --- | --- | --- |
| Zhang | 2017 | sPTB | term | maternal blood | genomics (GWAS) |
| Ackerman | 2015 | sPTB | term | placenta | transcriptomics (RNA-seq) |
| Heng | 2014 | sPTB | term | maternal blood | transcriptomics (microarray) |
| Chim | 2012 | sPTB | term | maternal blood | transcriptomics (microarray) |
| Mayor-Lynn | 2011 | sPTB | term | placenta | transcriptomics (microarray) |
| de Goede | 2017 | sPTB | term | cord blood | epigenomics (microarray) |
| Fernando | 2015 | sPTB | term | cord blood | epigenomics (microarray) |
| Parets | 2015 | sPTB | term | maternal blood | epigenomics (microarray) |
| Cruickshank | 2013 | sPTB | term | fetal blood | epigenomics (microarray) |
| Heng | 2015 | sPTB | term | maternal blood | proteomics (mass spec) |

### Data Transformation

Each of the 10 data sets was mapped to a gene-based format. This step was necessary because integRATE applies desirability functions both within and across studies and, in order for that integration to be possible, the genetic elements of each study have to match.

#### Genomics

SNP-based data containing P-values and effect sizes were mapped to genes with MAGMA, as outlined in the Zhang et al. supplementary methods (http://ctg.cncr.nl/software/magma) [35,47,48].

#### Transcriptomics

Gene expression data from microarray experiments were accessed via GEO (https://www.ncbi.nlm.nih.gov/geo/) and re-analyzed using the GEO2R plugin (https://www.ncbi.nlm.nih.gov/geo/info/geo2r.html [36-39]. Raw RNA-seq data from Ackerman et al. were analyzed in-house with custom scripts [33].

#### Epigenomics

Methylation data were accessed via GEO (https://www.ncbi.nlm.nih.gov/geo/) and re-analyzed using the GEO2R plugin (https://www.ncbi.nlm.nih.gov/geo/info/geo2r.html) [40-44].

#### Proteomics

Protein expression data were downloaded from supplementary files associated with each publication and the protein IDs were mapped to genes using Ensemble’s BioMart tool (https://www.ensembl.org/info/data/biomart/index.html) [34,45].

### Application of integRATE

After mapping results from all 10 omics studies to genes, we used integRATE to calculate desirabilities for all genes across all variables within studies. We ran four different sPTB analyses:

1. In the first analysis (**iR-none**), we ran integRATE with no added customizations (e.g., no cut points, no scales (i.e., scale coefficient = 1), no minimum or maximum desirabilities, etc.) (Fig. 3-5, Additional File 2).
2. In the second analysis (**iR-num**), we ran integRATE using numerical cut points (P = 0.0001, 0.1 and fold change = 1.5, 0.5, −0.5, −1.5) and no scales (Additional File 4-7).
3. In the third analysis (**iR-per**), we ran integRATE using percentile cut points (P = 5%, 95%, and fold change = 5%, 50%, 95%) and no scales (Additional File 8-11).
4. In the fourth analysis (**HardThresh**), we considered statistically significant genes from each study to represent the results that would have been obtained if the typical approach based on hard thresholds and intersection of significant genes across studies outlined earlier was applied (Additional File 12-13). All genes with adjusted P-values < 0.1 or unadjusted P-values < 0.05 were deemed significant in each study and intersected to compare with the results from integRATE [49].

To test whether the analyses described above produced results different from what might occur at random, we performed a permutation test shuffling desirabilities for all 26,868 genes 1,000 times.

**Fig. 3.**
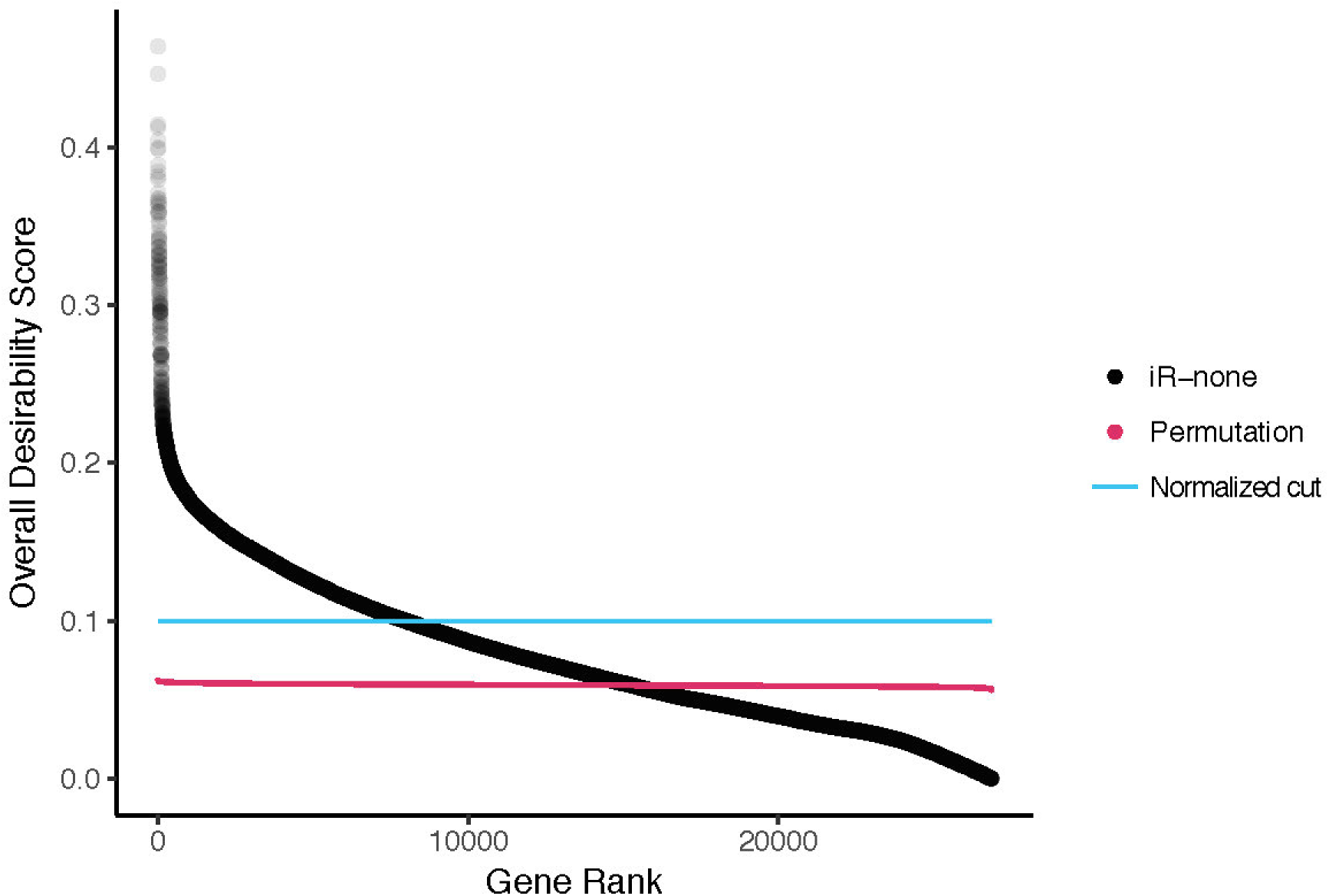
After integration, 7,977/26,868 genes were identified as highly desirable. All genes in the iR-none analysis were sorted from most desirable (rank = 1) to least desirable (rank = 26,868) and plotted according to their overall desirability scores, ranging from 8.04E-16 to 0.46. Because this analysis included 10 omics studies, the normalized lower bound for our set of ‘desirable’ candidate genes is 0.1 (blue line) and 7,977 genes achieved scores greater than or equal to that value. Furthermore, the results of our permutation test are plotted in pink, with a mean of 0.059 (95% CI [0.058, 0.061]). All desirability scores for the entire data set are available in Additional File 2 (and in Additional File 4 and 8 for iR-num and iR-per, respectively).

## Results

In total, our sPTB analyses integrated gene-based results from 10 omics studies (1 genomics, 4 transcriptomics, 4 epigenomics, and 1 proteomics; Table 1) and included data sets ranging from 422 genes [34] to 20,841 genes [38]. The null distribution generated by our random permutation test had mean desirability range from 0.056 to 0.062, with an average of 0.059 (95% CI [0.058, 0.061]) (Fig. 3).

### iR-none

First, the software was run without any added cuts, weights, or scales, resulting in a list of 26,868 genes with data from one or more of the 10 omics studies (Additional File 2). Normalized desirabilities for these 26,868 genes ranged from 8.04E-16 to 0.46 (mean = 0.08±0.05) (Fig. 3). Furthermore, 7,977 genes (29.7%) had desirabilities ≥ 0.1 corresponding to values equal to or higher than what would be achieved if a given gene achieved maximal desirability in one study but was absent from all others. These top 7,977 genes were enriched for 70 unique GO-Slim Biological Process categories, including pathways involved in metabolic processes, immunity, and signal transduction (Additional File 3) [50]. Additionally, 15,285/26,868 (56.9%) genes achieved desirabilities greater than the permutation mean of 0.059. The top 10 genes (Fig. 4 and 5) had desirabilities ranging from 0.46 (*CAPZB*) to 0.38 (*ACTN1*) and were all represented in each of the 10 omics data sets analyzed. This analysis applied integRATE without cut points, allowing for a straightforward, linear transformation of data across all variables and studies.

**Fig. 4.**
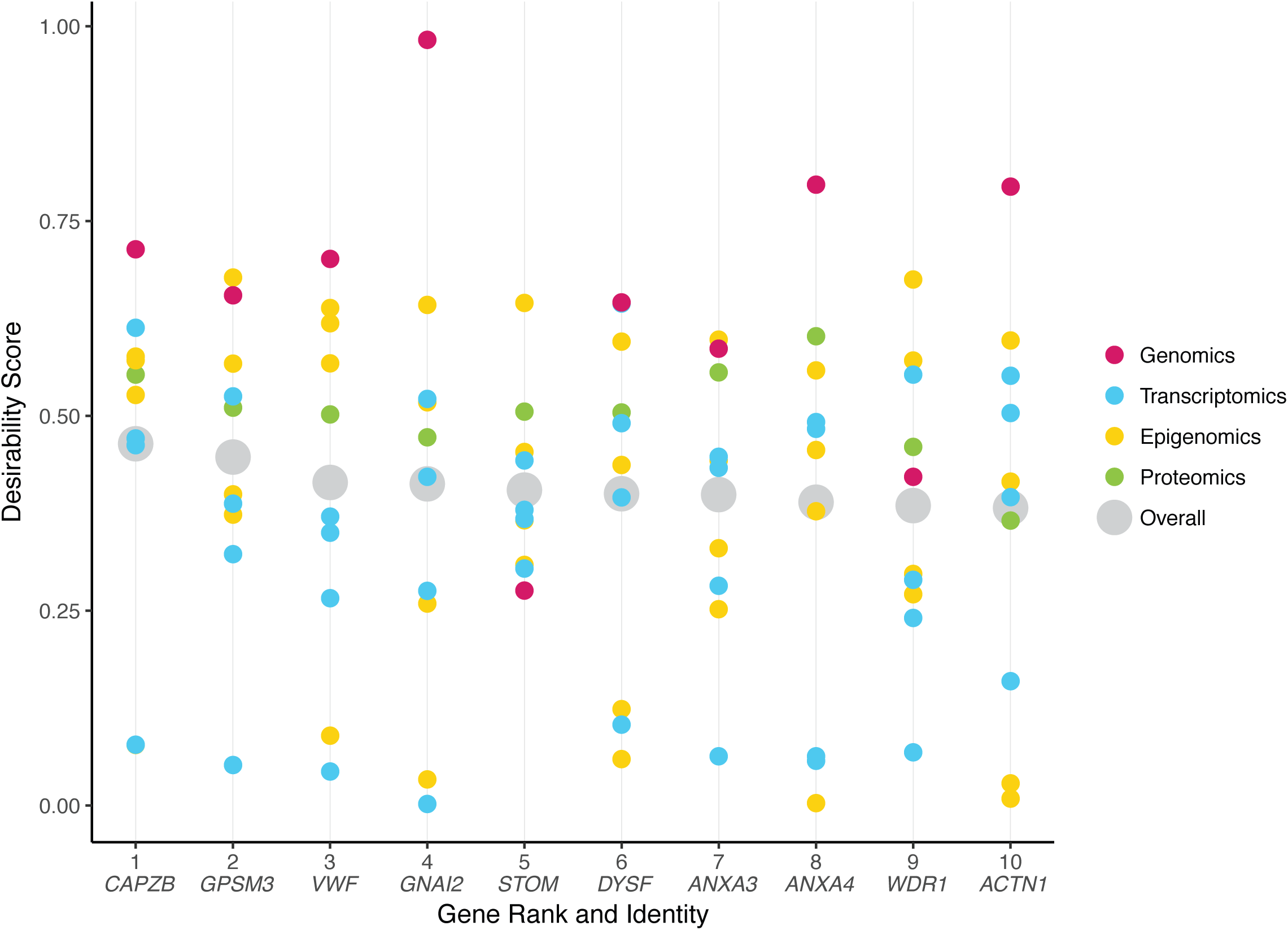
The top 10 most desirable genes have a wide range of desirabilities across data types. The top 10 genes in our analysis have overall desirabilities ranging from 0.38 (*ACTN1*) to 0.46 (*CAPZB*), but the *d_study_* values range, even when organized by data type. Some genes, like *STOM*, appear to be highly ranked not because of any extremely high *d_study_* value, but rather due to a lack of low *d_study_* values in any data type. In other words, this gene is likely not identified as particularly important in any individual study but shows a consensus of relatively strong evidence across all 10 studies. Contrastingly, other genes, like *CAPZB*, appear to be highly ranked due to one very high desirability score in a single data type (GWAS) that overpowers underwhelming evidence in other studies.

### iR-num

We next applied cut points based on numerical values (Additional File 4). P-values such that values smaller than 0.0001 received the maximum desirability score of 1 and values larger than 0.1 received the minimum desirability score of 0. All P-values between 0.0001 and 0.1 were transformed according to the *d_low_* function. For *d_extreme_* functions, 4 cut points were assigned and we chose commonly used values of 0.5 and 1.5 (or their equivalents if the values were log transformed). Therefore, fold changes below −1.5 or above −1.5 (or below log2(1/3) or above log2(3)) received the maximum desirability score of 1 and fold changes between −0.5 and 0.5 (or between log2(1/1.5) and log2(1.5)) received the minimum desirability score of 0. Intermediate values were transformed according to the *d_extreme_* function. This approach mirrors what was applied in a previous implementation of the desirability framework for omics data, and takes into account prior knowledge of typical P-value and fold change distributions [22]. While the top most desirable genes in iR-num appeared to be better candidates in each individual study (Additional File 7), using these cut points corresponding to standard significant P-value and fold change cut offs greatly reduced the number of desirable genes identified (Additional File 4). Specifically, only 1,386/26,868 (5.1%) genes achieved desirabilities greater than the permutation mean of 0.059 and the top 10 most desirable genes were analyzed by only 4 or 5 studies instead of all 10 (Additional File 6).

### iR-per

Finally, we applied cut points based on percentiles (Additional File 8). P-values were cut such that those in the top 5% received the maximum desirability score of 1 and those in the bottom 5% received the minimum desirability score of 0, with all values in between transformed according to the *d_low_* function. Fold changes were cut such that those in the top 5% and bottom 5% received the maximum desirability score of 1 and those in the middle 50% received the minimum desirability score of 0, with all other values transformed according to the *d_extreme_* function. In this analysis, 16,604/26,868 (61.8%) genes achieved desirabilities greater than the permutation mean of 0.059.

### HardThresh

For comparison, we also manually selected candidate genes by imposing a hard threshold on P-value (P-value < 0.05 if unadjusted and P-value < 0.1 if adjusted) (Additional File 12). After binning data into ‘significant’ gene lists, we intersected these lists to pull out genes that would have been identified simply by selecting the intersection of all significant genes. Although 18,727 genes were considered ‘significant’ in at least one study, no genes were identified as significant in all 10 studies. The top candidate gene (*KIAA0040*) was significant in 6/10 studies and 15 other genes were identified in 5/10 studies (Additional File 13). Interestingly, none of these 16 genes appear in the top 10 of our most desirable candidates after integration and, even more generally, none are specifically discussed in any of the studies, either.

### Using integRATE to identify the most desirable sPTB genes

In our sPTB pilot analyses, members of the annexin family (*ANXA3*, *ANXA4* and *ANXA9*) appear in the top 10 most desirable candidate gene sets regardless of analysis approach (e.g., without cut points as well as with numerical *and* percentile cut points). This family is involved in calcium-dependent phospholipid binding and membrane-related exocytotic and endocytotic events, including endosome aggregation mediation (*ANXA6*). In a previous proteomic analysis, *ANXA3* was found to be differentially expressed in cervicovaginal fluid 26-30 days before the eventual onset of sPTB as compared to before healthy, term deliveries [51]. Furthermore, members of the annexin family are known to be involved in coagulation (*ANXA3*, *ANXA4*). Coagulation has been previously suggested to be involved in PTB and, even though the mechanism of such involvement is still a mystery, it is interesting that several genes involved in coagulation or blood disorders appear in our top candidate lists [52]. In addition to *ANXA3* and *ANXA4*, *VWF* (or Von Willebrand Factor) is a gene encoding a glycoprotein involved in homeostasis that has been found to be expressed significantly more in preterm infant serum as compared to term [53,54]. Finally, another highly desirable candidate, *STOM*, encodes an integral membrane protein that localizes to red blood cells, the loss of which has been linked to anemia [55]. These results suggest that the development of sPTB may be linked to homeostasis pathways.

In addition to homeostasis and coagulation, another biological process represented across our results is actin regulation and muscle activity. The most notable gene associated with this biological process is *CAPZB*, which encodes part of an actin binding protein that regulates actin filament dynamics and stabilization and is present in the top 10 most desirable candidate gene list in all three analyses. Although *CAPZB* has never been linked to sPTB or other pregnancy pathologies, its role in muscle function could be linked to myometrial and uterine contractions that, when they occur prematurely, might be directly involved in the development of sPTB [56,57]. Another one of our top candidates, *ACTN1*, is also involved in actin regulation and, even more interestingly, has also been linked to blood and bleeding disorders [58,59]. Finally, several other highly desirable genes identified in one or more of our integrative analyses include *GPSM3*, *WDR1*, and *DYSF*, are all involved in the development and regulation of muscle or in the pathogenesis of muscle-related diseases [60-62].

Even outside the top 10 most desirable genes across our integrative analyses, we found genes both previously identified as being involved in pregnancy or sPTB pathology as well as involved in pathways potentially relevant to sPTB (Additional File 2). For example, one gene falling just outside the top 10 most desirable candidates in all analyses is *MMP9*, a matrix metalloproteinase. Interestingly, *MMP9* has been linked not only to sPTB, but also to PPROM and PE across a number of fetal and maternal tissues and at a variety of time points during pregnancy [63-67]. *MMP9* gene expression has been observed as significantly higher during preterm labor than during term labor in maternal serum, placenta, and fetal membranes [68-70]. Even in the first trimester, levels of *MMP9* in maternal serum were higher in PE cases than in healthy controls, suggesting that increased *MMP9* protein expression is linked to the underlying inflammatory processes governing PE pathogenesis [66]. Finally, fetal plasma *MMP9* concentration has been found to be significantly higher in fetuses with PPROM than in early and term deliveries with intact membranes, implicating *MMP9* in the membrane rupture mechanism controlling early delivery due to membrane rupture [67]. We see similar evidence of *MMP9* as a desirable sPTB candidate maintained across omics and tissue types in our integRATE analyses, raising the hypothesis that its role in inflammation and extracellular matrix organization relates to sPTB even in the absence of PPROM or PE.

## Discussion

By using desirability functions to rank genes within studies and combine results across studies, integRATE allows for the identification of candidate genes supported across experimental conditions and omics data types. This is especially important when heterogeneous sets of omics data, like those available for sPTB, where the statistical approaches developed for vertical or horizontal integration are challenging to apply. We have shown that integRATE can map any omics data to a common [0, 1] scale for linear integration and produce a list of the most desirable candidates according to their weight of evidence across available studies. These candidates then become promising targets for follow-up functional testing depending on where in the data their desirability signals come from. Analysis of 10 heterogeneous omics data sets on sPTB showed that the gene candidates identified using desirability functions appear to be much more broadly supported than those identified by the intersection of all significant genes across all studies and contain both genes that have been previously associated with sPTB as well as novel ones (Fig. 4 and 5, Additional File 13).

**Fig. 5.**
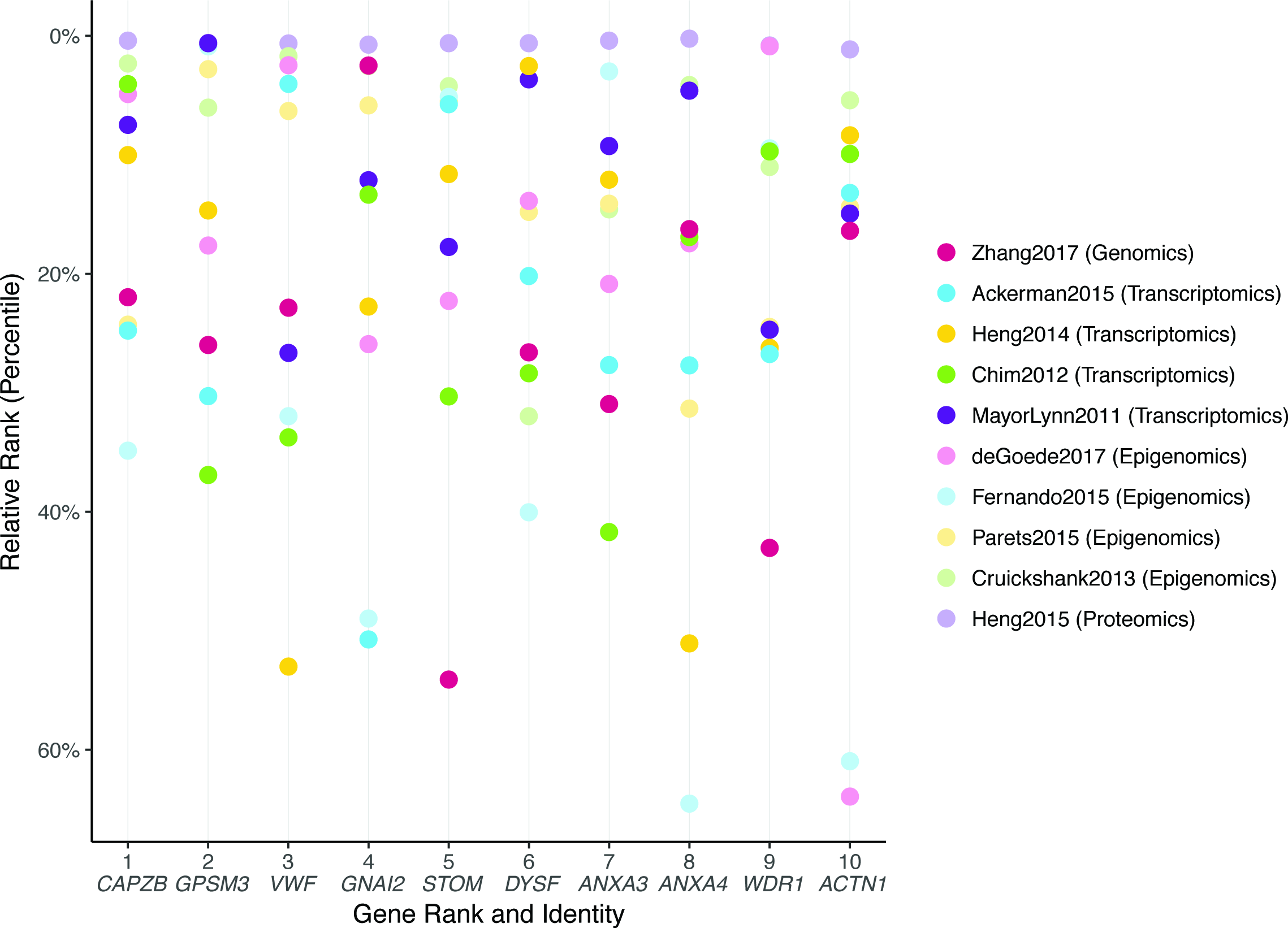
The top 10 most desirable genes also show a large discrepancy in their percentile ranks across studies. After ranking the genes in each study by desirability and calculating their percentiles based on the number of unique ranks, the top 10 most desirable genes appear to show even greater variability in relative ranking across not just data type, but individual studies. All 10 genes are in the top 25% of the (smaller) proteomics study, but their relative rankings vary significantly in all other studies. Furthermore, while none of the genes are in the top 25% of the GWAS study (Zhang 2017), other studies, like one of the transcriptomics analyses (Mayor-Lynn 2011), show a large range in relative rankings, with certain highly desirable genes ranked very high and others ranked very low.

integRATE identifies both known and novel candidate genes associated with a complex disease, including ones that are not be among the top candidates in any single omics study but are consistently (i.e., across studies) recovered as significantly (or nearly significantly) associated. For example, genes that are significantly differentially expressed at an intermediate to high level across *many* studies will have high desirability scores. Furthermore, integRATE can identify such genes across omics types, tissues, patient groups, and any other variable condition. Although integRATE allows for this kind of synergistic, desirability-based analysis, it is important to note that integRATE is not a statistical tool nor is it intended to be the end point of any analysis. Rather, it is a straightforward framework for the identification of well-supported candidate genes in any phenotype where true multi-omics data are unavailable and can also serve as a springboard for future functional analysis, an essential next-step in testing whether the candidates are actually involved in the biology of the disease or phenotype at hand.

Importantly, there is no single principled strategy for the selection of cut points. In our sPTB analyses (iR-none, iR-num, and iR-per), we observed that the imposition of cut points corresponding to generally agreed upon values (e.g., P-value < 0.0001) has the potential to greatly affect the resulting gene prioritization. On this basis, we propose that desirability functions are best used to integrate highly heterogeneous omics data without imposed numerical cut points for P-values, fold changes, and other variables. Implemented this way, one can maximize the information from the analysis of each omics data set used in prioritizing candidate genes. But users may also have reasons to want to put more weight on data sets that are of higher quality or on data types that may be more informative. In such instances, the weight parameter can be used to reflect study quality instead of imposing cut points (e.g., studies that fail to achieve P-values as low as others in the integrative analysis can be weighted less to reflect potentially lower experimental quality).

A recent GWAS analysis, the largest of its kind across pregnancy research, identified several candidate genes with SNPs linked to PTB [29]. This study linked *EBF1*, *EEFSEC*, and *AGTR2* to preterm birth and *EBF1*, *EEFSEC*, *AGTR2*, and *WNT4* to gestational duration (with *ADCY5* and *RAP2C* linked suggestively). By analyzing 43,568 women of European ancestry, this large study is the first to identify variants and genes that are statistically associated with sPTB. Interestingly, our integrative analysis identified *EBF1* as a desirable candidate (*d_overall_* = 0.15 [top 3%] in iR-none and *d_overall_* = 0.23 [top 1%] in iR-per), suggesting that this gene, in addition to GWAS, might also be functionally linked to sPTB pathogenesis across transcriptomics, epigenomics, and proteomics studies. Even when analyzing the 9 other omics studies without this GWAS data set, *EBF1* still achieved a *d_overall_* score of 0.17, placing it in the top 2% of all genes (Additional File 14). While our integrative analysis supports the identification of *EBF1* as an interesting candidate gene for follow up, the lack of signal for any of the other GWAS-identified hits also reinforces the need to approach complex phenotypes like sPTB from a variety of omics perspectives, since sequenced-based changes may impact the phenotype in indirect and complicated functional ways.

## Conclusions

Desirability-based data integration (and our integRATE software) is a solution most applicable in biological research areas where omics data is especially heterogeneous and sparse. Our approach combines information from all variables across all related studies to calculate the total weight of evidence for any given gene as a candidate involved in disease pathogenesis, for example. Although not a statistical approach, this method of data integration allows for the prioritization of candidate genes based on information from heterogeneous omics data even without known ‘gold standard’ genes to test against and can be used to inform more targeted downstream functional analyses.

## Abbreviations

*GWAS:*: genome-wide association study
*IUGR:*: intrauterine growth restriction
*PE:*: preeclampsia
*PPROM:*: premature rupture of membranes
*PTB:*: preterm birth
*sPTB:*: spontaneous preterm birth
*SNP:*: single nucleotide polymorphism

## Declarations

### Acknowledgements

We thank Dr. Lou Muglia for invaluable discussion and support in designing and applying this approach to data integration and Dr. Ge Zhang for providing access to preprocessed GWAS data.

### Funding

HRE was supported by a Transdisciplinary Scholar Award from the March of Dimes Prematurity Research Center Ohio Collaborative. This research was supported by the March of Dimes through the March of Dimes Prematurity Research Center Ohio Collaborative and the Burroughs Wellcome Fund.

### Availability of data and materials

All the data associated with and supporting the findings of this study are included in the manuscript and its supplementary files.

### Authors’ contributions

Conceived and designed experiments: HRE AR. Performed experiments: HRE. Developed scripts: HRE JS. Analyzed data: HRE. Wrote paper: HRE. Assisted with project development: AR JW PA JAC. Provided feedback: JW PA JAC AR.

### Ethics approval and consent to participate

Not applicable.

### Consent for publication

Not applicable.

### Competing interests

The authors declare that they have no competing interests.

## Supporting information

### Additional File 1. Results of meta-analysis to identify studies for integration

We outline the 10 studies meeting all inclusion criteria for integrative analysis. Furthermore, we list the other 44 studies that we identified through our literature search but we excluded from the data analysis as well as reasons for their exclusion.

### Additional File 2. All results from iR-none

All desirability scores across all variables in all studies as well as overall desirabilities and normalized overall desirabilities are presented.

### Additional File 3. GO-Slim gene set enrichment results

The PANTHER output for gene set functional enrichment is provided, including 37 statistically enriched biological pathways.

### Additional File 4. All results from iR-num

All desirability scores across all variables in all studies as well as overall desirabilities and normalized overall desirabilities are presented.

### Additional File 5. Results from iR-num

All genes in the analysis including numerical cut points were sorted from most desirable (rank = 1) to least desirable (rank = 26,869) and plotted according to their overall desirability scores.

### Additional File 6. Top 10 genes from iR-num by data type

Desirability scores for the top 10 most desirable genes are plotted according to the type of omics analysis.

### Additional File 7. Top 10 genes from iR-num by study

Desirability scores for the top 10 most desirable genes are plotted according to individual study.

### Additional File 8. All results from iR-per

All desirability scores across all variables in all studies as well as overall desirabilities and normalized overall desirabilities are presented.

### Additional File 9. Results from iR-per

All genes in the iR-per analysis were sorted from most desirable (rank = 1) to least desirable (rank = 26,869) and plotted according to their overall desirability scores.

### Additional File 10. Top 10 genes from iR-per by data type

Desirability scores for the top 10 most desirable genes are plotted according to the type of omics analysis.

### Additional File 11. Top 10 genes from iR-per by study

Desirability scores for the top 10 most desirable genes are plotted according to individual study.

### Additional File 12. Raw data for manual overlap based on significance dichotomization

All 18,727 genes identified as significant in at least 1 study and overlap across the entire data set.

### Additional File 13. Genes binned as significant in 4 or more omics studies

Upset plot showing intersections of significant genes across all 10 omics studies.

### Additional File 14. All results *without* Zhang 2017

All desirability scores across all variables in all studies as well as overall desirabilities and normalized overall desirabilities are presented.

